## Supplementary Information for "Smart sealants for prevention and monitoring of gastrointestinal anastomotic leaks using portable smartphone-controlled ultrasound transducers"

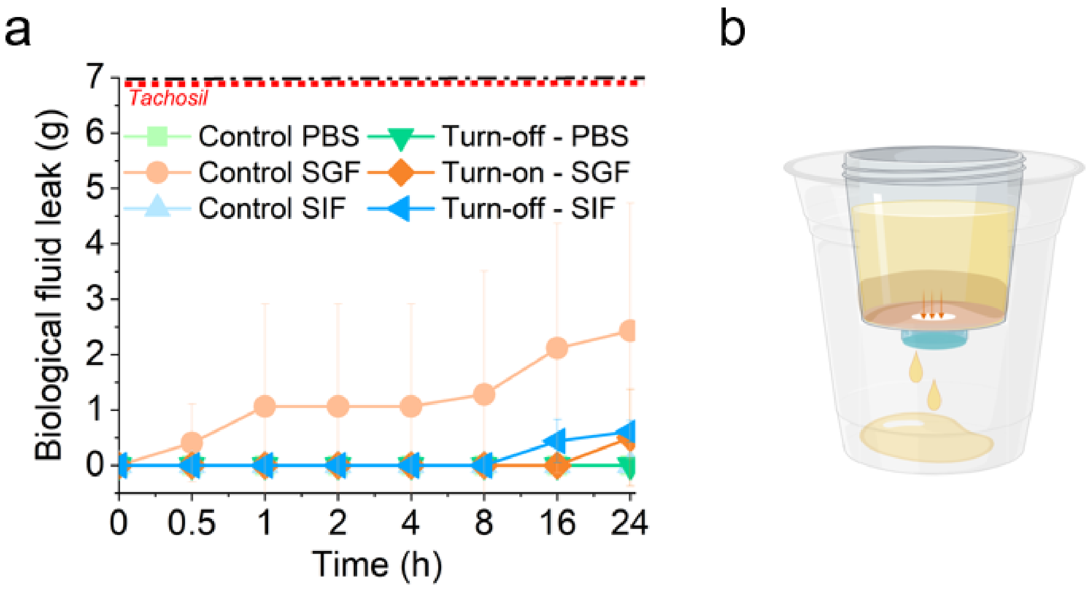


*Figure S1: **Intestinal fluid leakage model.** (a) Ex-vivo stationary model simulating the pooling of intestinal fluid and the sealing of a 4mm open hole in the tissue, covered by the sealant patch. Values for Tachosil included for reference. Leaked fluid mass of PBS, SIF and SGF, respectively, are monitored over time. Controls indicate tissue without hole.*

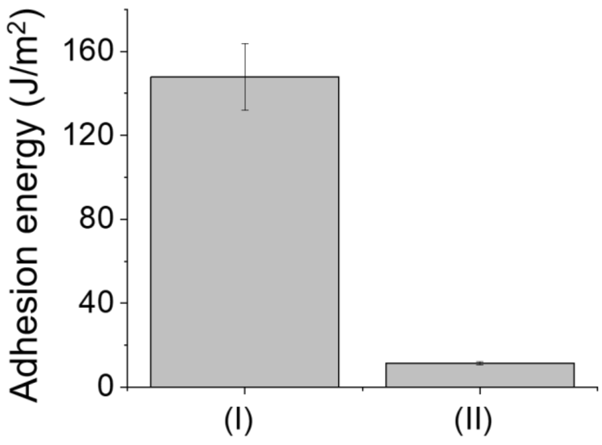


*Figure S2: **Adhesion energy with or without mIPN.** Adhesion energy (J/m$^2$) as a function of sample type in the T-peel setup of layered patches after application to porcine small intestine. (I) Layered patches consisting of a PAMPS support layer, a PNHEA backing, applied to tissue using a mutually interpenetrating network of PNAGA joining tissue and patch. (II) Layered patches consisting of a PAMPS support layer, a PNHEA backing and an interpenetrating network of PNAGA applied to tissue.*

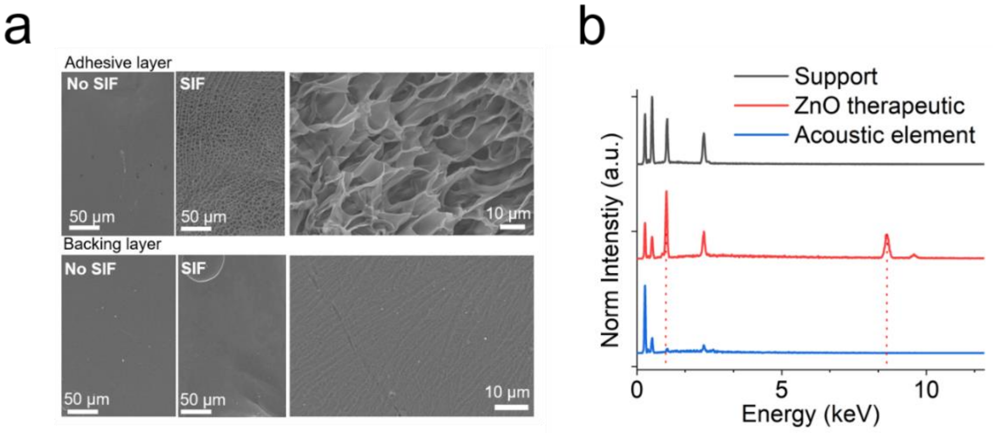


***Figure S3: Scanning electron micrographs of adhesive and backing layer before and after SIF exposure.*** *(a) SEM images of adhesive and backing layer following on sided contact with SIF, simulating a one sided contact with digestive fluid. (b) EDX elemental analysis of side-cut comprising ZnO therapeutic element, PAMPS support layer and Halo gas vesicle TurnOFF element. Red dotted lines denote the ZnLα and ZnKα peak locations of zinc.*

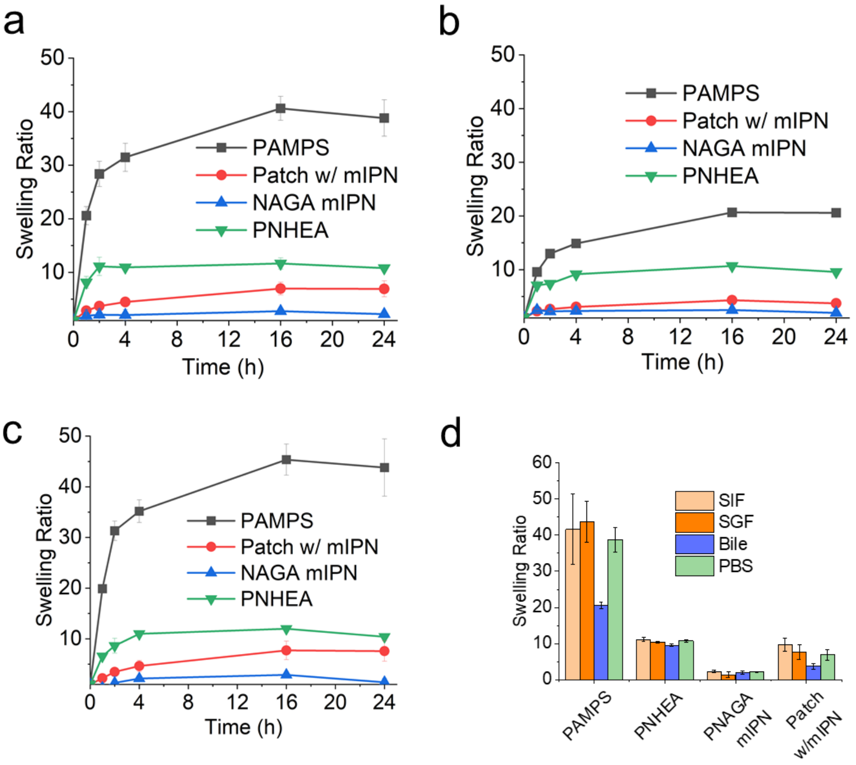


***Figure S4: Swelling behavior as a function of time and biological fluid.*** *Swelling of individual layers as well as fully assembled hydrogel patch after as a function of time in various simulated biological fluids at 37 °C.(a) Simulated gastric fluid, (b) Bile, (c)PBS, (d) 24 h swelling plateaus in different biological fluids.*

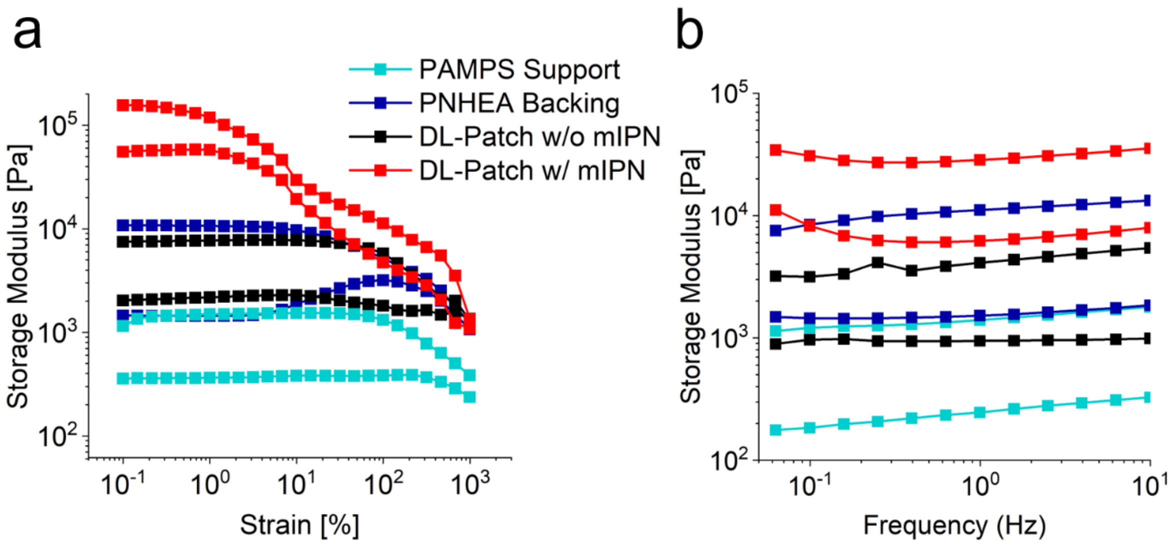


***Figure S5*: Rheological properties of hydrogel components and fully assembled DL-Patches.**

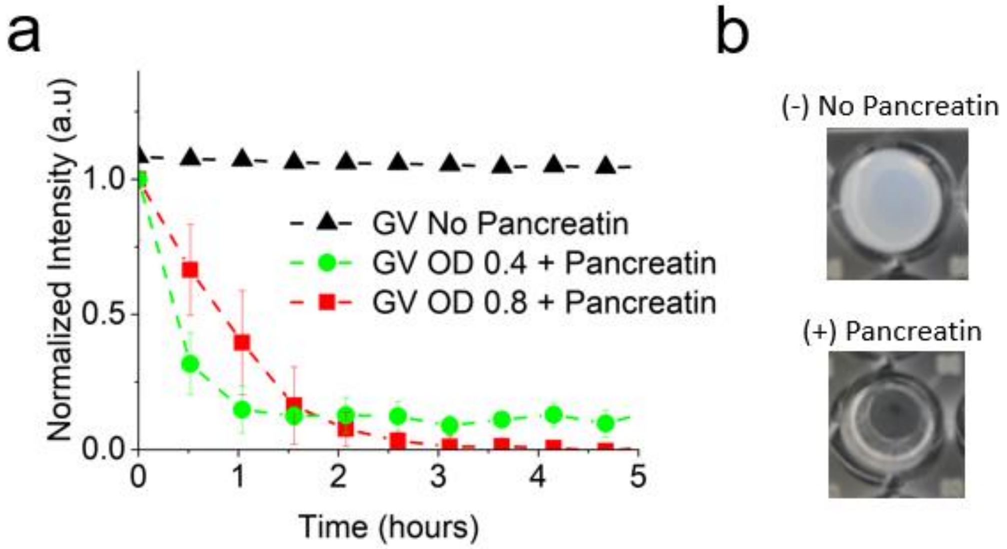


***Figure S6: Gas vesicle digestion kinetics.*** *(a) Digestion kinetics of gas vesicles following incubation with pocrine pancreatin enzyme. OD 0.4 corresponding to 20% vol solution in SIF salt background. (b) OD 0.8 in absence and presence of digestive enzyme.*

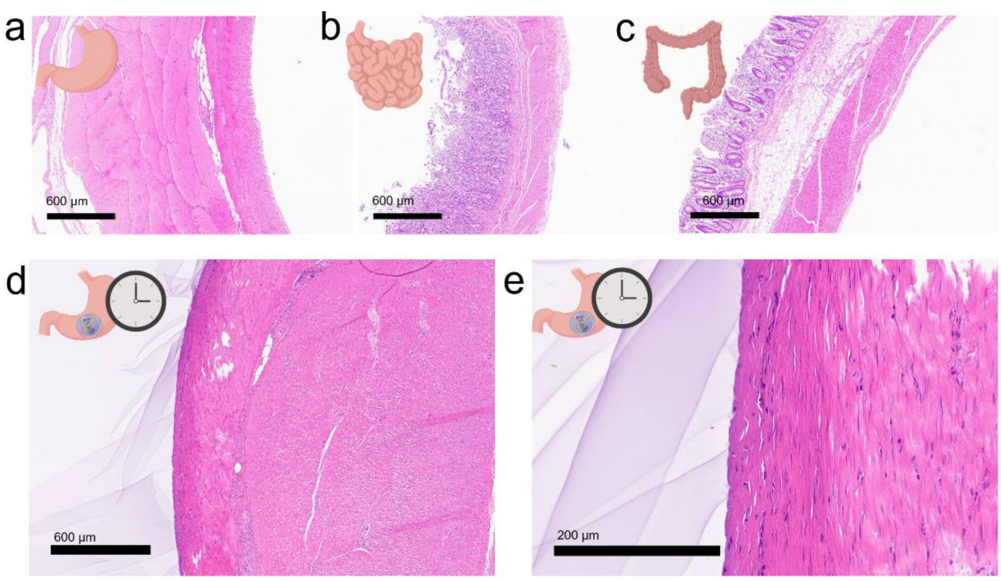


***Figure S7: Gastrointestinal tissue biopsies.*** *Attachment and interfacing of hydrogel sealant on various porcine tissues as showcased by H&E stained biopsy sections. (a –c) Bare tissue control samples showcasing native tissue serosa, (d-e) Representative images of tissue serosa with applied patch after 2h incubation with SGF.*

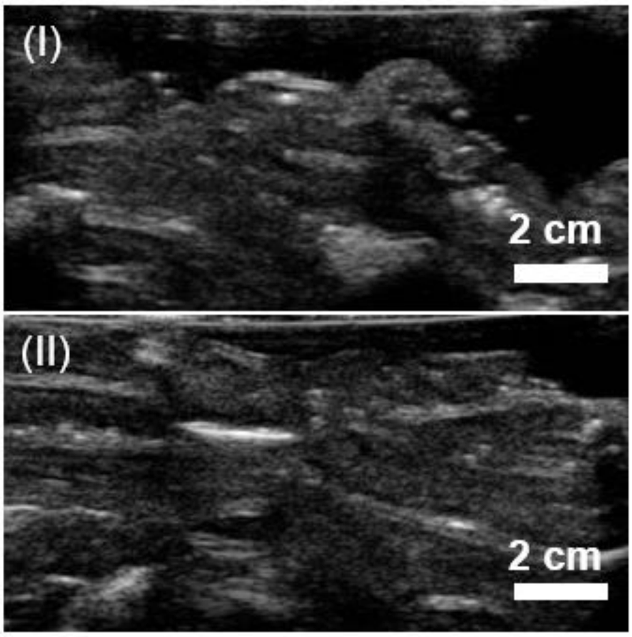


***Figure S8: Ultrasound imaging of DL-Patches on intestine.*** *Identification of gas vesicle loaded sensing element patch in an abdomen simulating model. (I) Native non-sealed porcine small intestine. (II) TurnOFF sealant patch attached on small porcine intestine.*

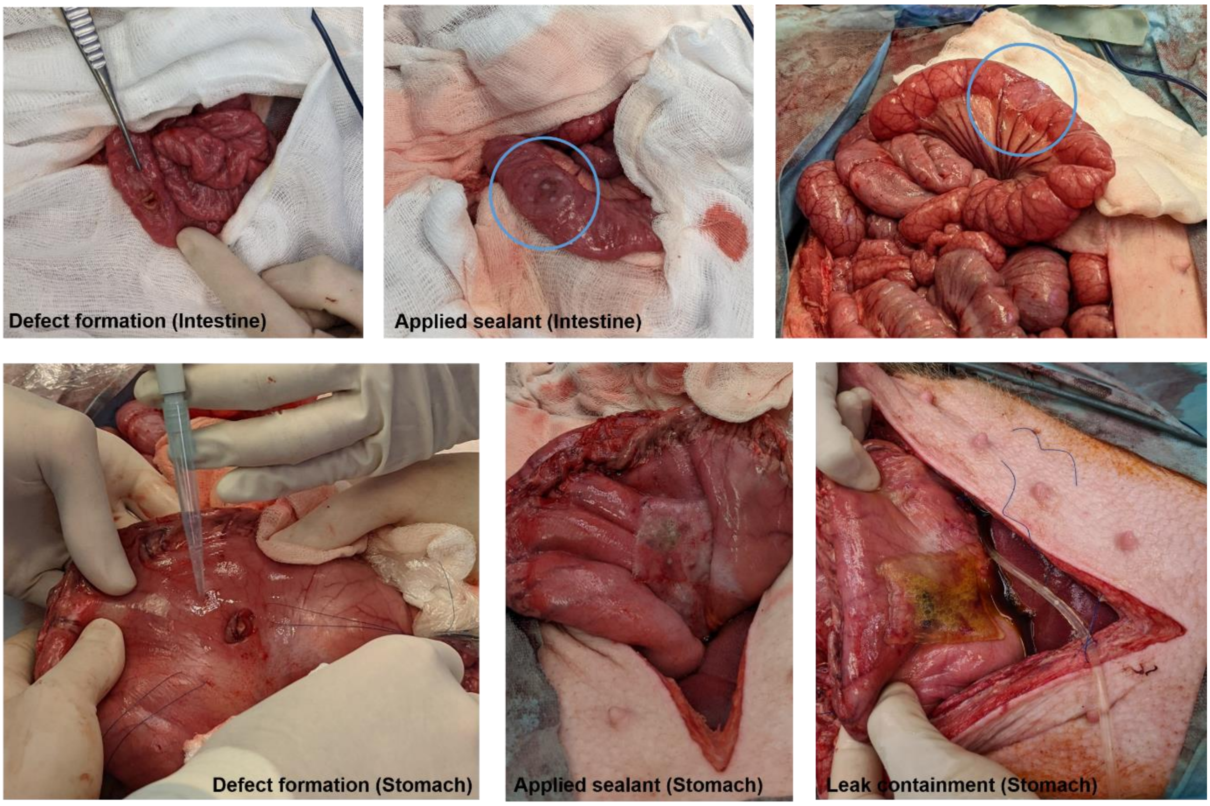


***Figure S9: Porcine model of gastrointestinal leaking.*** *Defect formation at the level of the small intestine and stomach, followed by DL-Patch application. Effective leak containment and firm patch adhesion even in presence of direct contact with SGF following a large defect (1x1 cm hole) on the stomach.*
